# Anti-oxidant and anti-inflammatory Effects of Aerosolised microalgal-derived extracellular vesicles in Bronchial Epithelial–Macrophage Co-cultures at the Air-Liquid Interface

**DOI:** 10.64898/2026.03.19.712886

**Authors:** Wesam Darwish, Giorgia Adamo, Mohammad Almasaleekh, Sabrina Picciotto, Paola Gargano, Daniele Paolo Romancino, Samuele Raccosta, Ralf Zimmermann, Mauro Manno, Antonella Bongiovanni, Sebastiano Di Bucchianico

## Abstract

Inflammation and oxidative stress are key drivers in the pathogenesis of chronic lung diseases, including asthma, pulmonary fibrosis, and chronic obstructive pulmonary disease. Extracellular vesicles derived from the marine microalga *Tetraselmis chuii*, referred to as nanoalgosomes, have recently gained attention as natural nanocarriers that possess inherent antioxidant and anti-inflammatory properties. In this study, we investigated the biocompatibility and protective effects of aerosolized nanoalgosomes in a bronchial epithelial–macrophage co-culture model at the air-liquid interface. Co-cultures of CALU-3 epithelial cells and differentiated THP-1 macrophages were primed with aerosolised nanoalgosomes and subsequently exposed to either oxidative stress (tert-butyl hydroperoxide) or an inflammatory stimulus (lipopolysaccharide; LPS). Epithelial barrier integrity and cytotoxicity were evaluated using transepithelial electrical resistance and lactate dehydrogenase release assays, respectively, while intracellular reactive oxygen species levels and cytokine secretion were measured to assess antioxidant and immunomodulatory responses. Nanoalgosomes were non-cytotoxic, preserved epithelial barrier integrity, and significantly reduced oxidative stress. In addition, nanoalgosomes priming attenuated LPS-induced secretion of pro-inflammatory cytokines (IL-1β, IL-6, IL-8, IL-18, TNF-α) as well as the anti-inflammatory cytokine IL-10, suggesting a balanced immunomodulatory response. Overall, aerosolized nanoalgosomes maintained epithelial homeostasis and mitigated both oxidative and inflammatory stress, underscoring their potential as a safe, sustainable, and effective therapeutic strategy for chronic inflammatory lung diseases. Given their natural origin, excellent biocompatibility, and suitability for aerosol delivery, nanoalgosomes represent a promising class of inhalable biotherapeutics.

## Introduction

Chronic lung diseases such as asthma, chronic obstructive pulmonary disease (COPD), and pulmonary fibrosis represent a major global health burden due to their prevalence, morbidity, and mortality. Central to the pathogenesis and progression of these diseases are persistent inflammation and oxidative stress [1]. These processes promote airway remodeling, leading to structural and functional alterations of the epithelial barrier and a reduction in pulmonary macrophage populations [2,3]. The complex interplay among cytokines, signaling pathways, and reactive oxygen species (ROS) contributes significantly to the physiological damage, ultimately exacerbating disease progression [4]. Despite their widespread clinical use, current therapeutics for respiratory diseases largely target symptom relief, providing short-term improvements in airflow or inflammation but failing to address the underlying cellular and molecular drivers of disease [4]. Consequently, there is an urgent need for strategies that simultaneously mitigate oxidative stress and inflammation to preserve epithelial barrier function and restore homeostasis in the lung microenvironment.

*Tetraselmis chuii* (*T. chuii*) is a non-toxic marine microalga, approved for human consumption as a novel food, and increasingly recognized for its rich content of bioactive compounds, including vitamins, carotenoids, polyphenols, phytosterols, superoxide dismutase (SOD), and polyunsaturated fatty acids (PUFAs) [5,6]. These compounds collectively contribute to the antioxidant and anti-inflammatory properties of *T. chuii*. Preclinical studies suggest that supplementation with *T. chuii* extracts can protect cells against oxidative damage induced by H₂O₂, increase enzymatic antioxidant expression, and reduce oxidative stress markers such as malondialdehyde in vivo [7–11]. Additionally, this alga modulates immune signaling by suppressing pro-inflammatory cytokines, including IFN-γ, TNF-α, and IL-1β, while regulating anti-inflammatory cytokines like IL-10 [7–9,12]. These multifaceted properties highlight the potential of *T. chuii* as a natural source of therapeutic biomolecules capable of restoring redox and inflammatory balance.

Extracellular vesicles (EVs) are lipid bilayer nanoparticles secreted by diverse cell types across species, expanding their relevance and potential for biomedical applications. They play a crucial role in intercellular communication by mediating the transport of biomolecules, including proteins, lipids, and nucleic acids. Their inherent biocompatibility and capacity for targeted delivery make them attractive candidates for novel biotherapeutics in chronic respiratory diseases [13–15]. EVs have been shown to alleviate inflammatory responses in the lung, reduce oxidative stress, and promote anti-inflammatory macrophage polarization, demonstrating their potential as both therapeutic agents and delivery systems [16–18].

Recently, naturally derived nanosized EVs extracted from *T. chuii* have been termed nanoalgosomes [19,20]. As *T. chuii* is rich in antioxidant and anti-inflammatory biomolecules, its EVs are likely to retain or convey these bioactivities while preserving the intrinsic structural and functional advantages of EVs, including biocompatibility, efficient cellular uptake, and protection of bioactive cargo from enzymatic degradation. Beyond their intrinsic bioactive properties, *T. chuii* offers a cost-effective, scalable, and sustainable source for EV production [21,22]. In vitro studies have shown that nanoalgosomes are non-toxic, decrease intracellular ROS levels, restore cytokine balance, and modulate antioxidant enzyme expression [19,20,22,23]. Using the in vivo model *Caenorhabditis elegans*, nanoalgosome supplementation was shown to significantly attenuate oxidative stress-induced locomotor decline [22]. The preservation of mobility provides compelling evidence for the systemic stability and protective effects of nanoalgosomes in a complex multicellular organism.

Despite these promising findings, the effects of nanoalgosomes in physiologically relevant airway models remain largely unexplored. To date, most studies have focused on simple monocultures, with only one study investigating pulmonary cells under submerged conditions [24]. This limits our understanding of how nanoalgosomes influence lung-specific responses in models that better recapitulate airway physiology [25]. Because oxidative stress and inflammation are interdependent drivers of lung pathophysiology, it is essential to determine whether nanoalgosomes can modulate intracellular ROS production and cytokine secretion within a more complex airway environment, including bronchial epithelial–macrophage interactions. Addressing this gap will provide critical insight into their potential to prevent or mitigate oxidative injury, immune dysregulation, and barrier dysfunction in chronic lung diseases such as asthma, COPD, and pulmonary fibrosis.

In this study, we evaluate the antioxidant and immunomodulatory properties of *T. chuii*-derived nanoalgosomes using a bronchial epithelial–macrophage co-culture model at the air-liquid interface (ALI). This physiologically relevant system enables assessment of airway barrier integrity, intracellular ROS modulation, and cytokine secretion in response to oxidative and inflammatory stimuli. By investigating the safety, biocompatibility, and functional effects of nanoalgosomes priming, we aim to determine their capacity to restore redox balance and regulate immune responses in the lung. Demonstrating that nanoalgosomes can reduce intracellular ROS, dampen pro-inflammatory cytokine production, and preserve epithelial barrier function would support their development as a biologically active, non-toxic, and scalable therapeutics for chronic respiratory diseases. Moreover, this work lays the groundwork for future translational efforts to leverage nanoalgosomes as inhalable delivery platforms, bridging natural bioactive compounds with next-generation medicine for pulmonary applications.

## Materials and methods

### Microalgae culture

*T. chuii* microalgae culture was performed following the established protocol described in Picciotto et al., 2025. Briefly, axenic *T. chuii* cultures were grown under sterile conditions in modified f/2 medium under controlled light (100 μmol m⁻² s⁻¹), temperature (22 °C ± 2 °C), and a 14:10 h light/dark cycle. Cell growth and viability were monitored weekly. At each time point, 1 mL of culture was sampled and optical density at 600 nm (OD₆₀₀) was measured spectrophotometrically. Cell concentration was calculated using a linear regression model (R² = 0.998) derived from a calibration curve correlating OD₆₀₀ values with cell counts obtained using a Thoma chamber from serially diluted cultures. 12 liters of culture were initiated at OD₆₀₀ = 0.300 and grown for 14 days to OD₆₀₀ = 0.600. Half of the culture volume (6 liters) was processed by tangential flow filtration (TFF) for nanoalgosome isolation, while the remaining half was replenished 1:1 with fresh f/2 medium and cultured for an additional two weeks before reprocessing.

### Tangential flow filtration

Upon reaching an optical density (OD₆₀₀) of approximately 0.600 after a two-week growth period, the cultures were processed. Nanoalgosome isolation was performed using TFF, a standard technique for purifying EVs [21]. All procedures were conducted under sterile conditions. The TFF system utilized a sequential three-stage filtration protocol with hollow fiber cartridges (Cytiva), progressing from a 650 nm pore size, to 200 nm, and finally to a 500 kDa molecular weight cut-off (MWCO). The system was equipped with in-line transducers and a peristaltic pump for continuous monitoring and control of permeate flow and transmembrane pressure, which were kept constant to ensure reproducible isolation conditions. During this process, the initial 650 nm microfiltration step removed intact cells, generating a permeate enriched with EVs. The subsequent 200 nm filtration retained larger vesicles while allowing the smaller nanoalgosomes to pass into the permeate. In the final stage, the 500 kDa UF membrane removed soluble proteins, small aggregates, and other non-vesicular contaminants, concentrating the nanoalgosomes in the retentate to a volume of ∼150 mL. A final concentration and diafiltration step was performed to exchange the native f/2 medium with calcium- and magnesium-free PBS, thereby further purifying and conditioning the nanoalgosome preparation. The final product was concentrated to an approximate volume of 7 mL.

### Nanoparticles tracking analysis (NTA)

The size distribution and concentration of three independent batches of nanoalgosomes was evaluated using the NanoSight PRO (Malvern Instruments), equipped with a blue (488 nm) laser and a high-sensitivity camera. Nanoalgosomes samples were diluted in filtered, particle-free water to achieve a particle concentration within the optimal detection range of 20–120 particles per frame, ensuring reliable measurements. Measurements were acquired using the Auto Camera and Auto Focus settings, with a flow rate of 3 µL/min. For each sample, five independent video captures were recorded and analyzed. Data were processed using NanoSight software version 2.0.

### Dynamic light scattering (DLS)

Nanoalgosomes samples (n=3) were diluted 1:5 and centrifuged at 1,000×g for 10 min at 4 °C to eliminate any dust particles. The supernatant was transferred to a quartz cuvette and incubated at 20 °C in the thermostatically controlled cell compartment of a BI200-SM goniometer (Brookhaven Instruments) equipped with a He-Ne laser (JDS Uniphase 1136P) tuned at 633 nm and a single-pixel photon counting module (Hamamatsu C11202-050). Scattered light intensity and its autocorrelation function g_2_(t) were measured simultaneously at a scattering angle θ = 90° using a BI-9000 correlator (Brookhaven Instruments). The electric field autocorrelation function *g_1_(t)* was calculated by using the Siegert relation:

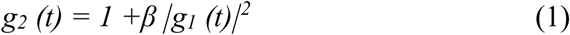

where *β* is an instrumental parameter and *g*_1_(*t*) is the field autocorrelation function, associated with the size (*σ*) of diffusing particles and their size distribution *P_q_(σ)* by the relation

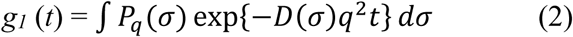

where *q = 4πn^λ^−1^sin(θ/2)* is the scattering vector in a medium with *n*^ refractive index and *D*(*σ*) is the diffusion coefficient of a particle of hydrodynamic diameter *D_h_ = σ*, determined by the Stokes−Einstein relation D(σ) = *k*_B_*T*[3*πησ*]^−1^, with *T* being the temperature, *η* the medium viscosity, and *k_B_* the Boltzmann constant. The size distribution *P*_q_(*σ*) was calculated under the assumption that the diffusion coefficient distribution is shaped as a Schultz distribution [21]. Two robust parameters are derived from this analysis: *D_z_*, the *z*-averaged hydrodynamic diameter, and *PDI*, the polydispersity index, an estimation of the distribution width. The field autocorrelation function of the samples was fitted with one component or two components if necessary

### Zeta-potential

Zeta potential measurements were performed using a Zetasizer Advance (Malvern Panalytical Ltd., UK). EV samples in PBS were suitably diluted in Milli-Q water to achieve comparable concentrations, while maintaining the same final dispersant (0.1× PBS). Samples were loaded into folded capillary cells, and measurements were carried out at 25 °C. Each measurement consisted of at least 12 automated runs, which were internally averaged by the instrument, was repeated three times (reported values correspond to the mean ± standard deviation). Zeta potential values were calculated from the measured electrophoretic mobility using the Smoluchowski approximation.

### Atomic force microscopy (AFM)

Sample preparation: glass slides were cleaned by boiling in acetone, dried by nitrogen stream and then exposed to UV radiation. The slides were then a) treated with a 0.25 M (3-aminopropyl)-triethoxysilane solution in chloroform for 3 minutes, rinsed with chloroform and dried with nitrogen; b) treated with 0.4 M glutaraldehyde aqueous solution for 3 minutes, rinsed with Milli-Q water and dried with nitrogen. 30 µl of vesicle solution were deposited onto functionalized glass slides at about 5 x 10^10^ particles/ml and incubated overnight. The samples were then gently rinsed with PBS to remove non-adsorbed vesicles. Vesicle imaging: Quantitative Imaging AFM measurements were performed in PBS using a Nanowizard III scanning probe microscope (JPK Instruments AG, Germany) equipped with a 15-μm z-range scanner, and AC40 (Bruker) silicon cantilevers (spring constant 0.1 N/m, typical tip radius 8 nm). The images (resolution 256×256 pixels) were acquired with a force setpoint of 130 pN and an extension speed of 25 μm/s. The cantilever was thermally calibrated by using the tool in JPK software [26].

### BCA assay and immunoblotting

The protein concentration of nanoalgosomes was quantified using the micro-bicinchoninic acid (BCA) Protein Assay Kit (Thermo Fisher Scientific). This colorimetric assay determines relative protein concentration by comparison to a bovine serum albumin (BSA) protein standard, from which a calibration curve was generated. The absorbance of the BCA soluble compound was measured at 562 nm using a GloMax Discover Microplate Reader. Protein separation was carried out by sodium dodecyl sulfate–polyacrylamide gel electrophoresis (SDS–PAGE) using 10% polyacrylamide gels. Equal amounts (∼5 μg) of cell lysates (*Tetraselimis chuii* and HEK-293) and nanoalgosome samples were mixed with appropriate volumes of 5× loading buffer (0.25 M Tris-HCl, pH 6.8, 10% SDS, 50% glycerol, 0.25 M dithiothreitol (DTT), and 0.25% bromophenol blue), heated at 100 °C for 5 min, and loaded onto the gels for electrophoretic separation. Cell lysates from *T. chuii* and Human Embryonic Kidney HEK-293 cells were included as internal reference controls to validate antibody specificity and to compare protein expression profiles, thereby supporting the identity of nanoalgosomes [27]. Following electrophoresis, proteins were transferred onto polyvinylidene difluoride (PVDF) membranes. Membranes were blocked for 1 h at room temperature in 3% BSA prepared in TBS-T (50 mM Tris-HCl, pH 8.0, 150 mM NaCl, 0.05% Tween 20). Primary antibodies against H⁺-ATPase (dil. 1:1000 in in 5% Milk/TBS-T1X, Agrisera), Alix (clone 3A9, dil. 1:150 in in 3% BSA/TBS-T1X, Santa Cruz), were incubated overnight at 4°C and 2 h at room temperature. After washing steps, membranes were incubated for 1 h with species-specific HRP-conjugated secondary antibodies (Cell Signaling). Blots were then washed thoroughly (four washes for 20 min, in TBS-T), and signal detection was carried out using the SuperSignal Pierce ECL (Thermo Fisher Scientific).

### DetectEV assay

To assess the membrane integrity and bioactivity of nanoalgosomes, the DetectEV assay [28] was used. This method quantify the intrinsic esterase-like activity within the lumen of EVs. The assay employs fluorescein diacetate (FDA), a lipophilic, non-fluorescent substrate that passively diffuses across intact lipid bilayers. Within intact vesicles, internal enzyme cleave FDA to produce fluorescein, a green-fluorescent molecule that accumulates in the lumen. For the assay, 2 × 10¹⁰ nanoalgosomes were added in triplicate to a 96-well plate and incubated with 18 µM FDA. The final reaction volume was adjusted to 200 μL with 0.2 µm-filtered PBS (without Ca²⁺ and Mg²⁺). Fluorescence (ex/em: 490/514 nm) was monitored continuously for 180 minutes using a GloMax Discover Microplate Reader (Promega). Specific enzymatic activity was quantified by comparison to a standard curve of fluorescein (0–300 nM). Activity is reported in units (U), defined as nM of fluorescein generated per minute (nM/min), calculated from the linear phase of the reaction and normalized to the total assay duration.

### Cell culture

A bronchial epithelial–macrophage co-culture model was established using the human bronchial epithelial cell line Calu-3 (ATCC, HTB-55) and differentiated macrophages derived from the human monocytic cell line THP-1 (ATCC, TIB-202) to investigate the anti-inflammatory and antioxidant properties of nanoalgosomes. Calu-3 cells were seeded onto 6-well Transwell inserts (Corning, CLS3450) at a density of 5 × 10⁵ cells per insert and maintained under submerged conditions in Minimum Essential Medium (MEM) GlutaMAX™ (Gibco, 41090093) supplemented with 10% (v/v) fetal bovine serum (FBS; Sigma-Aldrich, F7524), 1% (v/v) Penicillin–Streptomycin (Gibco, 15140122), and 1% (v/v) Non-Essential Amino Acids (Gibco, 11140050). Cultures were incubated at 37 °C in a humidified incubator containing 5% CO₂. The medium was replaced every 2–3 days, and the inserts were gently washed with pre-warmed Hank’s Balanced Salt Solution (HBSS; Gibco, 14025050) prior to medium change. After seven days of submerged culture, the apical medium was removed to establish ALI conditions. The apical surface was gently rinsed with pre-warmed HBSS the following day and every 2–3 days thereafter until exposure.

THP-1 cells were cultivated in DMEM/F-12 GlutaMAX™ medium (Gibco, 10565018) supplemented with 20% (v/v) FBS and 1% (v/v) Penicillin–Streptomycin under standard incubator conditions (37 °C, 5% CO₂). Differentiation into macrophages was induced by treating cells with 500 nM phorbol 12-myristate 13-acetate (PMA; Sigma-Aldrich, P1585) for 24 hours, followed by a 24-hour recovery period in fresh medium without PMA. One day prior to exposure, differentiated macrophages were detached using Accutase solution (Sigma-Aldrich, A6964) and 2.2 × 10⁵ cells were carefully seeded onto the apical surface of the CALU-3 epithelial layer. After 2-hour attachment period, the suspension medium was removed, and the co-cultures were maintained overnight under ALI conditions before subsequent treatments.

### ALI Exposure and Treatment Conditions

ALI exposures were performed on day 16 post-CALU-3 seeding using the VITROCELL® Cloud Alpha 6 system, which allows uniform and physiologically relevant aerosol deposition onto cultured cells. For nanoalgosome priming, 200 µL of a freshly prepared nanoalgosome suspension (ALG; 1.2 × 10¹³ EVs/mL) in calcium- and magnesium-free DPBS (Gibco, 14190144), spiked with 1% HBSS, was nebulized into the exposure chamber. The aerosol cloud was allowed to settle for approximately 5 minutes to ensure even deposition onto the apical surface of the epithelial–macrophage co-cultures. Following exposure, the inserts were incubated with fresh MEM medium without phenol red (Gibco, 51200038) in the basolateral compartment under standard culture conditions for 4 hours to permit cellular uptake and response to nanoalgosomes. To evaluate antioxidant activity, the apical compartment was treated with 1 mL of 1 mM tert-butyl hydroperoxide (TBHP; Thermo Fisher Scientific, A13926) in MEM medium without phenol red for 4 or 24 hours to induce oxidative stress. For assessment of anti-inflammatory effects, 200 µL of lipopolysaccharide solution (LPS; Santa Cruz Biotechnology, sc-221855) in ultrapure water at 125 or 250 µg/mL was nebulized onto the apical surface of the co-culture, followed by a 24-hour incubation at 37 °C. Solvent controls (SCs) were prepared in parallel with all treatment conditions. First, all SC inserts were nebulized with 200 µL of calcium- and magnesium-free DPBS supplemented with 1% HBSS and subsequently incubated for 4 hours. For oxidative-stress experiments, 1 mL of MEM medium without phenol red was then added to the apical compartment and inserts were incubated for 4 or 24 hours. For inflammation studies, SC inserts were instead nebulized with 200 µL of sterile ultrapure water containing 1% HBSS and incubated for 24 hours at 37 °C. An incubator control (IC) was included by maintaining co-cultures at the ALI under standard culture conditions with fresh basolateral medium only.

Based on published data, the VITROCELL® Cloud Alpha 6 system provides a consistent aerosol deposition efficiency of approximately 15.6% ± 0.6%, ensuring uniform and reproducible delivery of nebulized materials across different experimental conditions [29]. Accordingly, the estimated deposition for nanoalgosome priming is 1.34 × 10¹⁰ EVs/cm², while for LPS concentrations of 125 and 250 µg/mL, the corresponding deposited doses are approximately 140 and 280 ng/cm^2^, respectively.

### Cytotoxicity

At each experimental endpoint, transwell inserts were gently washed with 1 mL of pre-warmed HBSS to collect residual lactate dehydrogenase (LDH). Both the apical wash and basolateral medium were then collected and centrifuged at 250 × g for 5 minutes. Cytotoxicity under all experimental conditions was subsequently evaluated using the Cytotoxicity Detection Kit^PLUS^ (Roche, 4744926001) according to the manufacturer’s instructions. LDH release, which serves as a reliable indicator of membrane damage and overall cytotoxicity, was quantified by transferring 50 µL of each sample in duplicate into a 96-well plate, followed by the addition of 50 µL of the LDH reaction mixture. The plates were incubated at room temperature for 20 minutes to allow for color development, after which the optical density was measured at 492 nm and 620 nm using a Varioskan LUX Multimode Microplate Reader. As a positive control (PC), one insert was treated with 2% (v/v) Triton X-100 (Sigma-Aldrich, T8787) for 20 minutes prior to the end of the exposure. The combined optical density of the apical and basolateral fractions was then used to calculate cytotoxicity as a percentage relative to the PC. All measurements were performed in three independent experiments, and each condition was analyzed in duplicate to ensure reproducibility and statistical reliability.

### Transepithelial Electrical Resistance

At each experimental time point, the integrity of the CALU-3/dTHP-1 layer was assessed using an EVOM™ Epithelial Volt/Ohm Meter 3 (World Precision Instruments) equipped with 4 mm wide chopstick electrodes. This method was employed to monitor the tight junction formation and barrier function of the bronchial epithelial monolayer under different treatment conditions. Prior to the measurements, 1 mL of pre-warmed HBSS was added to both the apical and basolateral compartments of each Transwell insert to maintain electrical continuity across the membrane. The electrodes were carefully inserted, and stabilized transepithelial electrical resistance (TEER) values were recorded in triplicate from three separate points on each insert to obtain representative measurements from different regions of the insert. To normalize the TEER measurements, the resistance of a cell-free insert was subtracted from each recorded value. The corrected resistance was then multiplied by the surface area of the insert to obtain TEER values expressed in Ω·cm². The mean of the resulting data was calculated from four independent experiments and used as an indicator of epithelial barrier integrity and viability following exposure to oxidative or inflammatory stimuli.

### Intracellular reactive oxygen species

The 2ʹ,7ʹ-dichlorodihydrofluorescein diacetate (H_2_DCFDA; Sigma-Aldrich, 287810) assay was used to evaluate the anti-oxidative effects of nanoalgosomes in CALU-3/THP-1 co-cultures. Prior to nanoalgosome priming, transwell inserts were incubated with 50 µM H_2_DCFDA in MEM without phenol red (Gibco, 51200038) for 1 hour at 37 °C to allow intracellular uptake. Following this preloading step, the inserts were exposed to nanoalgosomes and treated with TBHP as previously described to induce oxidative stress. After the treatment period, the inserts were washed, and the cells were detached using 0.25% Trypsin-EDTA (Gibco, 15400054) for 3 minutes. The cell suspensions were collected and centrifuged at 200 × g for 7 minutes to pellet the cells. The resulting pellets were resuspended in DPBS and transferred to a black 96-well plate for fluorescence measurement at 488 nm excitation and 530 nm emission every 5 minutes. ROS levels were calculated as relative fluorescence units and expressed relative to control samples. Measurements were taken within the linear range of the signal (0–20 minutes), with data acquired from three technical replicates per condition across three independent experiments.

### IL-8 and IL-18 quantification

The levels of the pro-inflammatory cytokine IL-8 and IL-18 were measured using the Human IL-8/CXCL8 DuoSet ELISA Kit (R&D Systems, DY208) and Human Total IL-18 DuoSet ELISA (R&D Systems, DY318) following the manufacturer’s recommendations. Both kits follow identical experimental procedures. Briefly, a 96-well plate was coated with 100 µL of capture antibody and incubated overnight at room temperature. Plates were then washed three times with wash buffer consisting of 0.05% Tween-20 (Sigma-Aldrich, P9416) in DPBS (Gibco, 14190144). Wells were blocked for 1 hour with 300 µL of blocking buffer containing 10 mg/mL BSA (Sigma-Aldrich, A2153) in DPBS. Following an additional wash step, 100 µL of apical washes or standards were added to the wells and incubated for 2 hours. Standards were prepared as serial two-fold dilutions, ranging from 31.2 to 2000 pg/mL for IL-8 and from 11.7 to 750 pg/mL for IL-18. Plates were washed three times before adding 100 µL of detection antibody and incubating for an additional 2 hours. Following a fourth wash, 100 µL of streptavidin–horseradish peroxidase was added and incubated for 20 minutes. After a final wash, 100 µL of 3,3′,5,5′-tetramethylbenzidine (TMB; Sigma-Aldrich, T0440) substrate was added and incubated for 20 minutes. The enzymatic reaction was stopped by adding 50 µL of 2 N H₂SO₄ (Fisher Chemical, S/9240/PB15), and absorbance was measured at 450 nm with a reference wavelength of 570 nm.

### Luminescent ELISA

Luminescence-based ELISA was used to quantify the levels of IL-1β, IL-6, IL-10, and TNF-α (Promega; W6010, W6030, W6070, W6050) according to the manufacturer’s guidelines. In a white 96-well plate, 50 µL of apical washes were added in duplicate alongside seven standards, with standard curves ranging from 18.2 pg/mL to 25,000 pg/mL prepared using 3.33-fold serial dilutions. Subsequently, 50 µL of freshly prepared antibody mixture was added to each well, and the plate contents were mixed at 300 rpm for 10 seconds. Plates were then incubated at 37 °C in a CO₂ incubator for 60 minutes. Following incubation, the plate was equilibrated to room temperature, and 25 µL of luminescence detection reagent was added. The plate was briefly shaken at 400 rpm for 10 seconds and incubated for 4 minutes before measuring luminescence using a Varioskan LUX Multimode microplate reader (Thermo Scientific). For IL-1β specifically, 50 µL of each sample was assayed with seven standards ranging from 21.7 pg/mL to 40,000 pg/mL, prepared by 3.5-fold serial dilutions. The plate was also incubated for 75 minutes instead of 60 minutes, following the manufacturer’s recommendations.

### Statistical analysis

All assays were conducted in at least three independent experiments. Data are presented as the mean ± standard error of the mean (SEM). Statistical significance was determined using one-way analysis of variance (ANOVA) with Bonferroni multiple comparison test. Graphs and data analyses were generated using GraphPad Prism for Windows (version 10.2.3; GraphPad Software).

## Results

### Production and quality assessment of EV derived from the microalga *Tetraselmis chuii*

To generate a reproducible source of nanoalgosomes, an axenic culture of *T. chuii* was successfully cultivated under strictly controlled sterile conditions. At harvest, *T. chuii* culture were processed for nanoalgosome isolation using TFF. This optimized preparative process achieved over a 1000-fold enrichment of the small extracellular vesicle fraction, yielding 5 mL of purified small EVs in PBS from the initial 6 L culture volume. After TFF-based isolation, nanoalgosomes were systematically characterized using standardized quality assessment assays, as recommended by the MISEV2023 guidelines and reported in the literature [22,28,30,31] (Figure 1).

**Figure 1.**
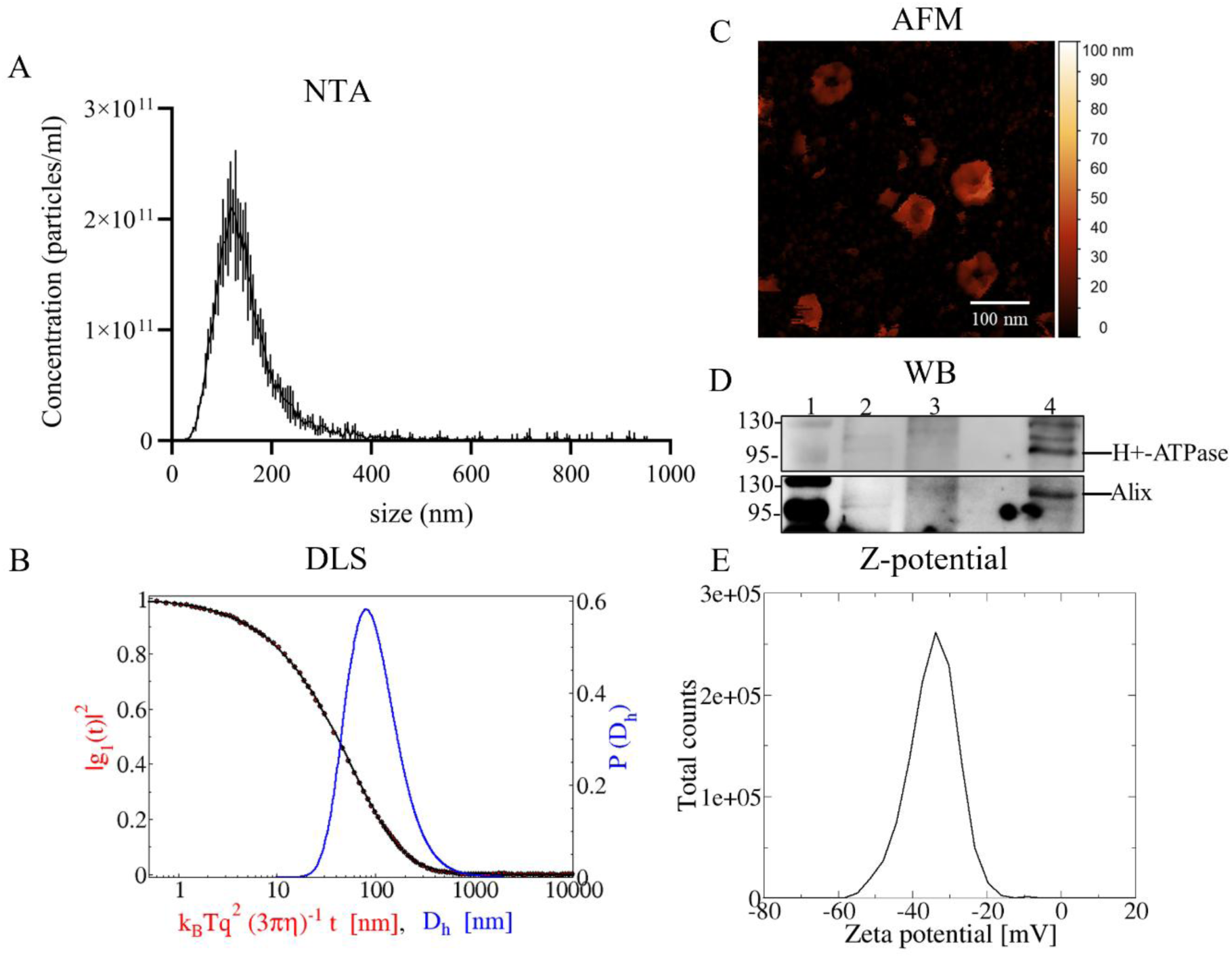
Nanoalgosome features. (**A**) concentration and size distribution were determined by NTA (error bars represent five measurements per sample) and (**B**) confirmed by DLS; (**C**) morphology was assessed by AFM; (**D**) expression of canonical EV markers (H⁺-ATPase and Alix) was analyzed by immunoblot in the following samples: 1) marker; 2) HEK-293 lysate; 3) *T. chuii* lysate; 4) nanoalgosomes; (**E**) nanoalgosome surface charge was measured as zeta potential. Representative curves and images for all analyses are shown for three independent samples.

Analysis of nanoalgosomes preparations (n = 3) by BCA revealed a protein content of 148 ± 9 µg/mL. NTA in scattering mode showed a particle concentration of 5.6 × 10¹² ± 8.7 × 10¹¹ particles/mL, as well as a size distribution characterized by a modal size of 123 ± 3 nm (Figure 1A). The distribution percentiles were D10 = 80 ± 4 nm, D50 = 137 ± 3 nm, and D90 = 253 ± 5 nm, indicating that 10%, 50%, and 90% of the particles were below these diameters, respectively. The NTA profile also suggested a narrow size distribution, consistent with a population of small EVs. This was confirmed by DLS measurements, which showed hydrodynamic diameters (Dz) ranging from 105 ± 5 nm to 123 ± 5 nm with polydispersity indices (PDI) between 0.47 ± 0.01 and 0.49 ± 0.01 (Figure 1B).

AFM further supported these results, confirming the characteristic small EV-like morphology of nanoalgosomes (Figure 1C). In addition, protein analysis by SDS-PAGE and immunoblotting detected canonical nanoalgosome biomarkers, including H⁺-ATPase and Alix proteins (Figure 1D) [19]. Furthermore, nanoalgosomes dispersed in PBS exhibited a zeta potential value of −30 ± 4 mV, confirming the expectation of a negative surface charge, as for EVs from other cell sources (Figure 1E). Overall, these findings demonstrated that nanoalgosomes derived from *T. chuii* exhibit the characteristic size, morphology, surface charge, and molecular markers of small EVs, with consistently low batch-to-batch variability.

Moreover, to verify the functionality of the nanoalgosomes batches used in this study, the DetectEV assay was performed. The DetectEV assay quantify the enzymatic activity of intraluminal esterase-like enzymes, that is contextually a measure of vesicle membrane integrity. The results showed a bioactivity equal to 1 ± 0.1 nM/min in nanoalgosomes samples used in this study (n = 3). This value is fully consistent with the bioactivity range reported for high-quality nanoalgosomes preparations [28], confirming the reproducibility of the production process and the suitability of the batches used for subsequent cell experiments.

### Cytotoxicity assessment of the CALU-3/THP-1 Co-culture

Cytotoxicity was assessed by measuring LDH release as an indicator of plasma membrane damage, which results in the leakage of the cytoplasmic LDH enzyme. Pro-oxidant stimulation with TBHP caused a modest increase in LDH release after 4 hours of exposure, an effect that was attenuated when cells were pre-treated with nanoalgosomes, which on their own did not affect cell viability (Figure 2A). No significant cytotoxicity was detected after 24 hours (Figure 2B). LPS nebulization at 125 and 250 µg/mL, corresponding to a deposition of approximately 140 and 280 ng/cm^2^, resulted in an increase in cytotoxicity after 24 hours, with the effect at 250 µg/mL reaching statistical significance compared with SC (Figure 2C). Notably, nanoalgosomes priming did not enhance cytotoxicity and instead restored LPS-induced changes to control levels.

**Figure 2.**
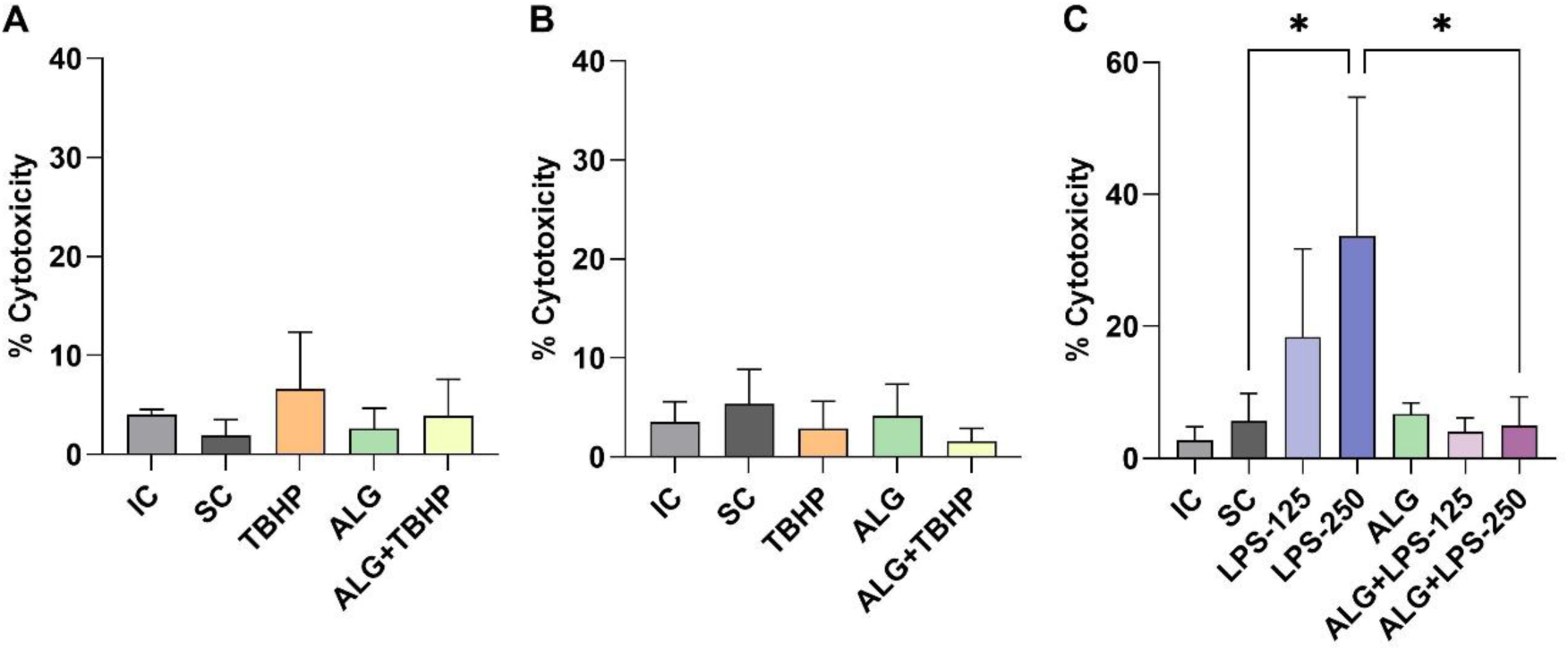
Cytotoxicity of co-cultured CALU-3 and dTHP-1 cells as determined by LDH release following oxidative stimuli (1mM TBHP) for 4 hours (**A**), 24 hours (**B**), or following inflammatory stimuli (125 and 250 µg/mL LPS) for 24 hours (**C**). Data are expressed as mean ± SEM of at least three independent experiments. One-way ANOVA was used to determine statistically significant differences among the tested groups. (*p < 0.05).

Overall, neither nanoalgosome priming nor subsequent oxidative stimuli elicited significant cytotoxic effects, in contrast to the more pronounced cytotoxicity observed under inflammatory challenge. These findings highlight the safety and biocompatibility of nanoalgosomes, and support their potential suitability for airway-targeted therapeutic applications.

### Integrity of the Bronchial Epithelial Barrier at the Air-Liquid Interface

Epithelial barrier integrity was evaluated by measuring the TEER in CALU-3/dTHP-1 co-cultures exposed for 4- or 24 hours to SC, TBHP, nanoalgosomes. or TBHP following nanoalgosome priming. TEER values in SC treated co-cultures remained stable at both time points, indicating intact tight junctions and functional epithelial barrier. Priming with aerosolized nanoalgosomes did not significantly alter TEER, confirming their biocompatibility and absence of disruption of airway epithelial integrity (Figure 3A, B). Exposure to 1 mM TBHP for 4- or 24 hours resulted in TEER values similar to SC, with only a modest, non-significant reduction observed in TBHP-treated cultures (SC: 1537 ± 119.5 Ω·cm² vs. TBHP: 1401 ± 90.5 Ω·cm²) (Figure 3A). This indicates that oxidative stress at this concentration did not compromise epithelial barrier function, either acutely or after extended exposure. Nebulization of LPS at 125 µg/mL and 250 µg/mL for 24 hours did not significantly affect TEER compared to SC, with only a slight increase observed at the highest concentration (Figure 3C). Notably, this minor effect was absent in co-cultures primed with nanoalgosomes. Collectively, these results demonstrate that nanoalgosomes preserve epithelial barrier integrity under both oxidative or inflammatory challenges, further supporting their biocompatibility.

**Figure 3.**
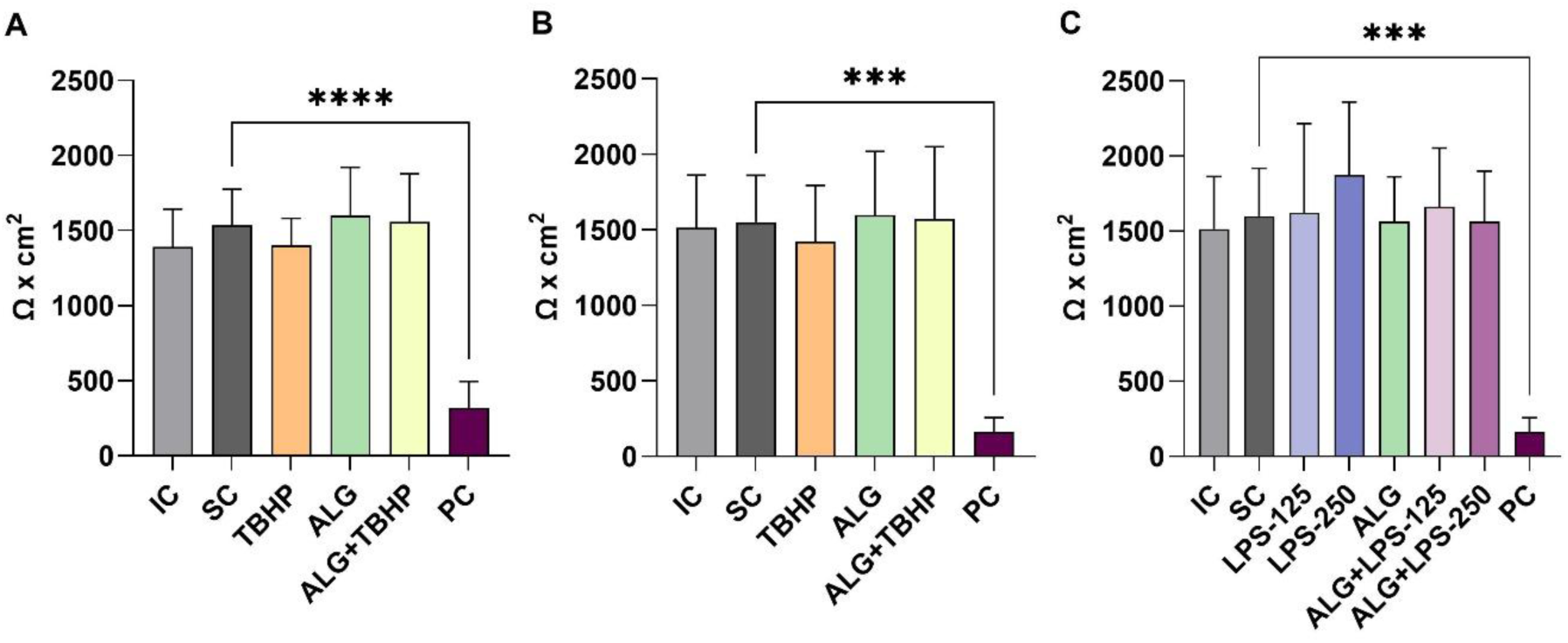
Effect of nanoalgosome priming on epithelial barrier integrity in CALU-3/dTHP-1 coculture. TEER values after 1mM TBHP treatment for 4 hours (**A**) and 24 hours (**B**), and in response to nebulized LPS (125 and 250 µg/mL) after 24 hours (**C**). Data are presented as mean ± SEM of four independent experiments. Statistical significance was determined using one-way ANOVA (**p < 0.01, ****p < 0.0001).

### Effects of Nanoalgosome Priming on Intracellular ROS Production Following Oxidative Stimuli

The antioxidant activity of nanoalgosomes was evaluated by measuring intracellular ROS in CALU-3/dTHP-1 co-cultures using the H_2_DCFDA assay. Nanoalgosomes treatment alone did not affect basal ROS levels (Figure 4A, B), indicating that the vesicles are non-oxidant with both macrophages and bronchial epithelial cells [22]. As expected, exposure to 1 mM TBHP for 4 hours induced a robust oxidative response, increasing intracellular ROS by approximately 4.8-fold relative to the SC. Notably, 4 hours of nanoalgosome priming markedly mitigated this response, reducing TBHP-induced ROS accumulation by 46% (Figure 4A). Although ROS levels in the ALG+TBHP group remained above control values, the reduction reflects a substantial protective effect against acute oxidative stress. At 24 hours post-treatment, ROS levels in TBHP-exposed cultures declined toward baseline (Figure 4B), consistent with endogenous antioxidant defense mechanisms over time. Even at this later time point, nanoalgosomes-primed cultures exhibited approximately 55% lower ROS levels than TBHP-treated cells, demonstrating a sustained antioxidant effect. While the magnitude of protection differed between the 4- and 24 hours measurements, the overall trend indicates that nanoalgosomes confer both early and prolonged resistance to oxidative stress.

**Figure 4.**
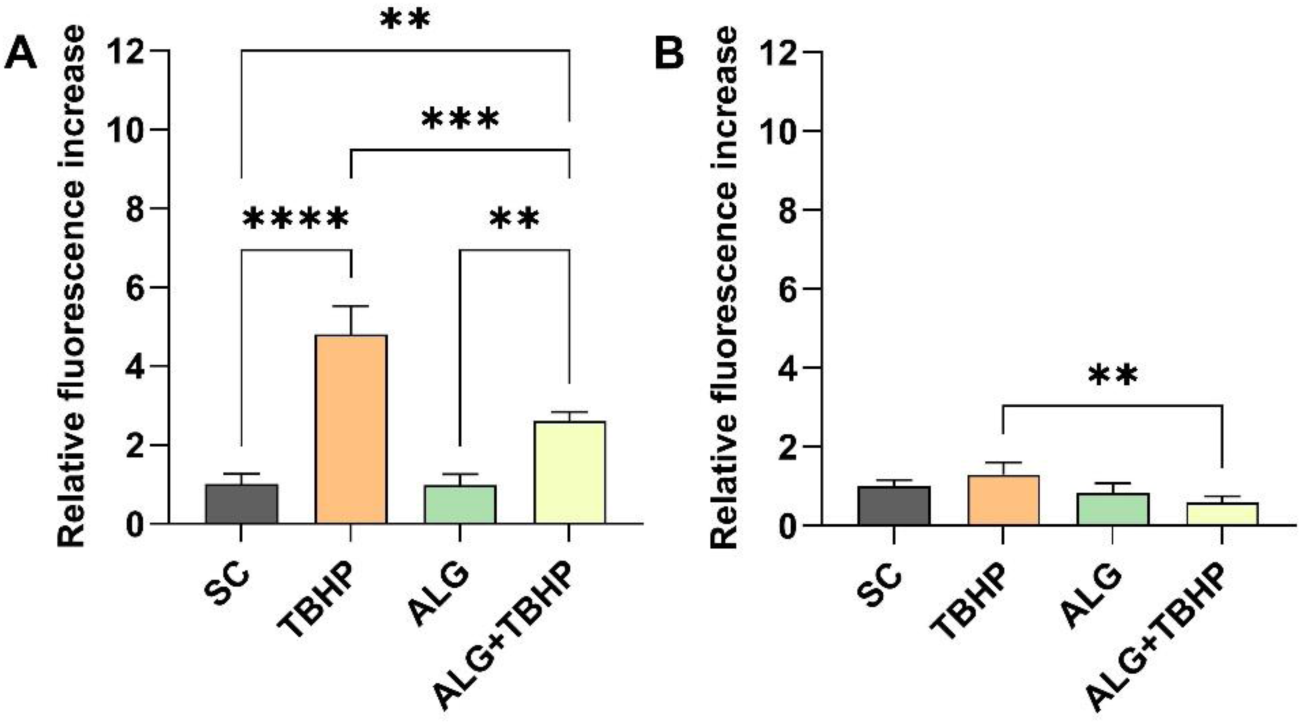
Intracellular ROS levels following TBHP treatment. Intracellular ROS generation was quantified using the H_2_DCFDA assay and expressed as relative fluorescence intensity after exposure to 1 mM TBHP for 4 hours (**A**) and 24 hours (**B**). Data are presented as mean ± SEM from four independent experiments. Statistical significance was determined using one-way ANOVA (**p < 0.01, ***p < 0.001, ****p < 0.0001).

Taken together, these findings indicate that nanoalgosome pre-exposure significantly mitigates intracellular ROS generation in response to TBHP-induced oxidative stress. This supports the intrinsic antioxidant capacity of nanoalgosomes and their potential to preserve redox balance in airway epithelial systems.

### Effects of Nanoalgosome Priming on Cytokine Secretion Induced by LPS Nebulization

To investigate the immunomodulatory properties of nanoalgosomes, cytokine secretion was quantified 24 hours after exposure to nebulized LPS (125 and 250 µg/m). Nanoalgosome priming alone did not significantly alter the secretion of any of the six cytokines measured relative to SCs, indicating that nanoalgosomes do not disrupt basal immune signalling (Figure 5).

**Figure 5.**
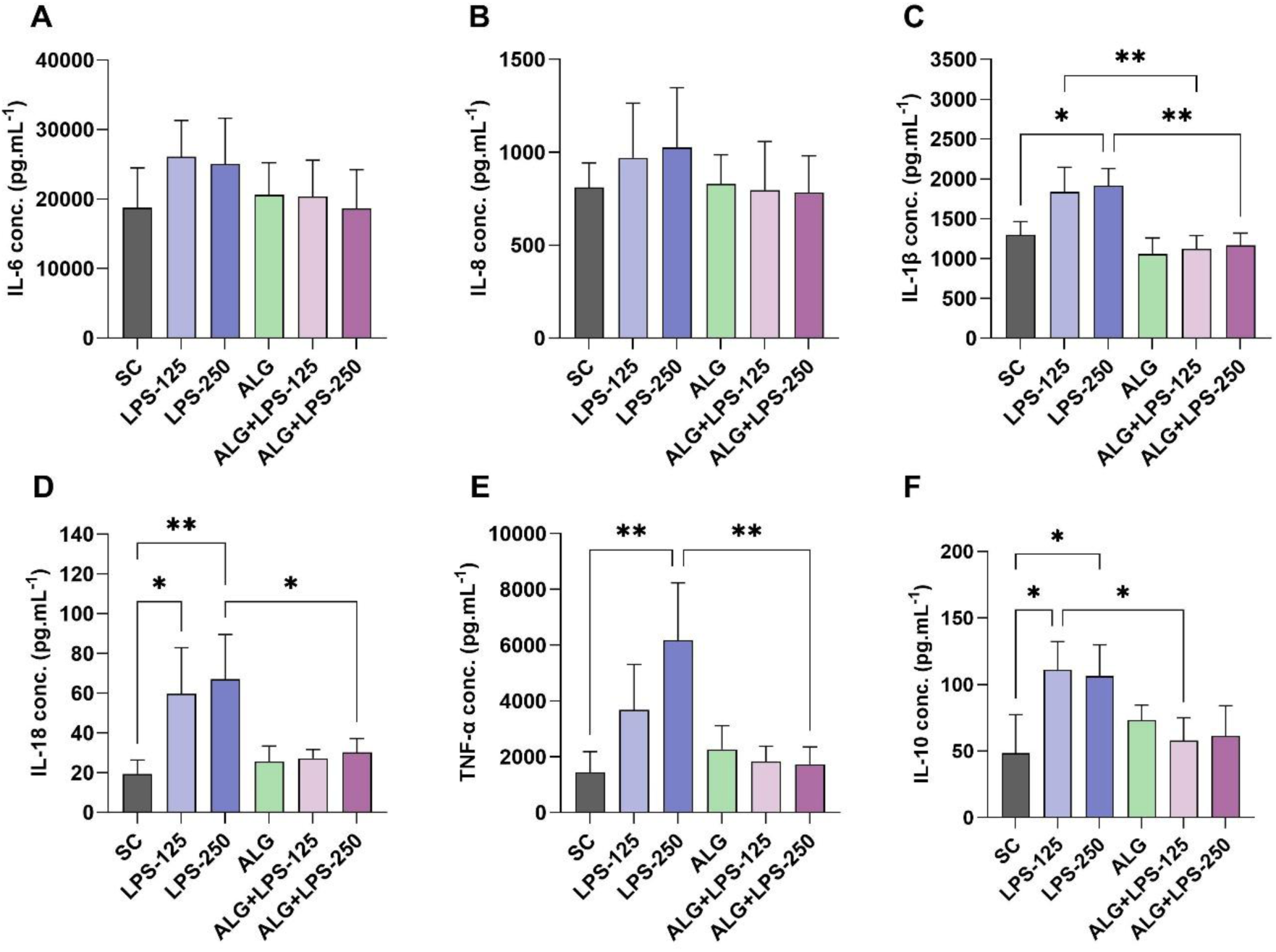
Cytokine secretion profile of CALU-3/dTHP-1 coculture following exposure to nebulized LPS. (**A**) IL-6, (**B**) IL-8, (**C**) IL-1β, (**D**) IL-18, **(E)** TNF-α, and (**F**) IL-10 were quantified after exposure to 125 and 250 µg/mL nebulized LPS solution for 24 hours. Data are presented as mean ± SEM from three independent experiments. Statistical significance was determined using one-way ANOVA (*p< 0.5, **p < 0.01).

IL-6 levels remained statistically unchanged across all conditions, showing only a slight upward trend (Figure 5A). A similar pattern was observed for IL-8, where modest LPS-induced increases were attenuated following nanoalgosome pre-treatment (Figure 5B). In contrast, more robust pro-inflammatory responses were detected for IL-1β, IL-18 and TNF-α: both concentrations of LPS elevated their secretion, confirming activation of the inflammatory pathway in the co-culture (Figures 5C-E). Nanoalgosome priming effectively mitigated these increases. At the higher LPS concentration (250 µg/mL), IL-1β, IL-18 and TNF-α levels were reduced by approximately1.6-fold, 2.2-fold and 3.6-fold, respectively, relative to LPS alone. The anti-inflammatory cytokine IL-10 was also significantly elevated following LPS exposure (Figure 5F). Notably, nanoalgosome priming restored IL-10 secretion toward baseline, reducing LPS-induced elevations by 1.9 and 1.7-fold at 125 µg/mL and 250 µg/mL LPS, respectively.

Overall, these findings indicate that nanoalgosomes exert a modulatory effect on both pro- and anti-inflammatory cytokine response. By attenuating both pro- and anti-inflammatory cytokine dysregulation, nanoalgosomes seem to promote a more balanced cytokine milieu, likely through the modulation of macrophage-epithelial interactions and the restoration of airway inflammatory homeostasis.

## Discussion

This study demonstrates that EVs derived from *Tetraselmis chuii*, referred to as nanoalgosomes, are biocompatible and exert protective antioxidant and anti-inflammatory effects in a physiologically relevant bronchial epithelial–macrophage co-culture model. Using CALU-3 epithelial cells and PMA-differentiated THP-1 macrophages at the ALI, we showed that nanoalgosome priming did not compromise cell viability or epithelial barrier integrity, which are essential parameters for preventing pathogen invasion and progressive lung damage. These findings confirm the non-cytotoxic nature of nanoalgosomes and their compatibility with airway cellular systems. Similar observations have been observed in vitro in diverse cell lines, as well as in *C. elegans* and mice, where nanoalgosomes did not compromise cell or organism viability [19,32]. Likewise, *T. chuii* extracts have been shown to preserve endothelial barrier function in human brain microvascular endothelial cells, reinforcing the notion that *T. chuii*-derived biomaterials are non-disruptive to epithelial and endothelial structures regardless of their form [11].

The maintenance of epithelial integrity following nanoalgosome exposure may relate to the regulation of tight junction proteins, including E-cadherin, which is essential for epithelial cohesion and proliferation, in addition to the production of pro-inflammatory mediators and growth factors [33]. Recent evidence indicates that nanoalgosomes upregulate E-cadherin mRNA expression, suggesting a potential mechanism by which they may actively reinforce epithelial barrier stability [24]. This feature is particularly relevant for chronic lung diseases such as COPD, asthma, and pulmonary fibrosis, in which epithelial dysfunction and loss of barrier integrity are key pathogenic events.

Exposure to TBHP induced a pronounced increase in intracellular ROS after 4 hours, consistent with oxidative stress-mediated injury commonly reported in airway epithelial cells exposed to pollutants or inflammatory mediators [34,35]. The lack of ROS elevation at 24 hours likely reflects the activation of endogenous antioxidant defense systems over time, as ROS generation is typically an early and transient event [36,37]. Remarkably, nanoalgosome priming significantly attenuated this oxidative response, suggesting an intrinsic antioxidant capacity capable of mitigating oxidative insults. This protective effect likely arises from the bioactive cargo naturally present in *T. chuii*, including carotenoids, polyphenols, and PUFAs which are known to enhance cellular antioxidant defenses [5,6].

These antioxidant mechanisms may involve Nrf2 activation, upregulation of enzymatic defenses such as SOD, catalase, and glutathione peroxidase, and mitigation of lipid peroxidation, as reflected by reduction in malondialdehyde levels [7–9,38,10]. Consistent with this, nanoalgosomes have been shown to modulate oxidative stress–related enzymes including AKR1C2 (aldo-keto reductase family 1 member C2), FTH1 (ferritin heavy chain 1), Alox12 (arachidonate 12-Lipoxygenase), CAT (catalase), GPX1 (glutathione peroxidase 1) and GSR (glutathione reductase) in mammary epithelial cells [22]. In alveolar epithelial cells, nanoalgosomes also restored glutathione peroxidase 4 levels, further supporting their role in redox regulation [24]. Notably, the magnitude of intracellular ROS reduction observed here was comparable to that achieved using liposomes encapsulating drugs such as pirfenidone or quercetin—agents known to enhance treatment outcomes in lung fibrosis and non-small cell lung cancer [24,39–41]. Together, these findings suggest that natural nanoalgosomes can function as biologically active nanocarriers with intrinsic antioxidant properties.

The cytokine profiling further highlights the immunomodulatory potential of nanoalgosomes. LPS nebulization induced a typical pro-inflammatory response characterized by elevated IL- 1β, IL-6, IL-8, IL-18 and TNF-α secretion, reflecting macrophage and epithelial activation [42–49]. The minimal increase in IL-6 and IL-8 may be attributed to apical LPS exposure, since Toll-Like Receptor 4, the primary receptor for LPS in bronchial epithelial cells [50], is typically expressed at low levels and predominantly localized on the basolateral side of the cells [51]. High concentrations of pro-inflammatory cytokines are characteristic of airway inflammatory conditions such as COPD, asthma, and pulmonary fibrosis [52–59]. Nanoalgosome priming markedly reduced the secretion of these cytokines, particularly IL-1β, IL-18, and TNF-α, indicating a robust anti-inflammatory response. This attenuation is likely mediated by the inhibition of NF-κB and MAPK signaling cascades, pathways that are known to be modulated by the polyphenolic and fatty acid components of *T. chuii* [6]. In macrophage cultures, nanoalgosomes have been shown to downregulate IL-6 expression, which aligns with our findings in a more complex epithelial–macrophage system [22]. Importantly, by normalizing TNF-α release, nanoalgosomes may help preserve epithelial tight junction integrity, since excessive TNF-α is known to disrupt barrier function and contribute to airway hyperresponsiveness and fibrosis [60].

IL-10 is a key anti-inflammatory cytokine released by both macrophages and bronchial epithelial cells in the lung [61,62]. Interestingly, IL-10 levels were elevated following LPS exposure but normalized upon nanoalgosome priming. This normalization suggests that nanoalgosomes facilitate a return to immune homeostasis by preventing excessive compensatory anti-inflammatory signaling. Given that IL-10 dysregulation is associated with chronic inflammation, COPD, and cancer, the ability of nanoalgosomes to stabilize IL-10 levels highlights their potential in rebalancing immune responses in diseased airways [63–65].

The anti-inflammatory properties of nanoalgosomes likely stem from their distinctive bioactive composition. *T. chuii* is rich in vitamins, carotenoids, polyphenols, phytosterols, and PUFAs, which may act synergistically to modulate inflammatory signaling pathways [66]. These compounds have been shown to suppress the production of key pro-inflammatory cytokines, including IFN-γ, TNF-α, and IL-1β [7–9,12]. Recent studies further demonstrated that *T. chuii*-derived nanoalgosomes downregulate the expression of transforming growth factor-β (TGF-β) in alveolar epithelial cells, highlighting their potential to mitigate inflammation at the cellular level [24]. Given that cytokine modulation is an emerging therapeutic strategy for managing inflammatory lung diseases, nanoalgosomes may represent a multifaceted alternative to conventional cytokine-blocking therapies, offering a natural means of attenuating excessive cytokine responses [67–69].

From the perspective of therapeutic development, the natural origin, nanoscale size, and aerosolization compatibility of nanoalgosomes render them particularly suited for inhalation delivery. Aerosolized administration allows for targeted deposition in the respiratory tract, maximizing local bioavailability and minimizing systemic side effects—a major advantage over conventional systemic anti-inflammatory or antioxidant drugs. Moreover, their innate antioxidant and anti-inflammatory properties could complement or enhance the therapeutic performance of encapsulated drugs, offering opportunities for combination strategies in targeted pulmonary delivery. Notably, the effects of native nanoalgosomes were comparable to those of liposomes loaded with established drugs such as pirfenidone and quercetin, further highlighting their therapeutic promise even in the absence of additional cargo [24].

Further research is warranted to decode the molecular cargo of nanoalgosomes, including proteins, lipids, and microRNAs, which likely orchestrate their protective actions through modulating cellular signaling pathways related to oxidative stress response, inflammation, and tissue repair. Additional studies employing in vivo animal inhalation models will be essential to validate the therapeutic efficacy, optimize dosing regimens, and evaluate safety profiles and pharmacokinetics under physiologically relevant conditions.

In summary, *T. chuii*-derived nanoalgosomes represent a new class of naturally sourced EVs capable of modulating oxidative and inflammatory processes in airway systems at the ALI. Their intrinsic bioactivity, combined with safety and scalability, highlights their potential as both therapeutic agents and delivery vehicles in the treatment of chronic lung diseases characterized by oxidative stress and inflammation, including asthma, COPD, and pulmonary fibrosis.

## Conclusion

This study demonstrates that *Tetraselmis chuii*–derived nanoalgosomes are biologically active EVs following aerosolization with potent antioxidant and anti-inflammatory properties relevant to chronic lung diseases characterized by epithelial barrier disruption, oxidative stress and immune dysregulation. Nanoalgosome priming effectively protected bronchial epithelial–macrophage co-cultures at the ALI by reducing intracellular ROS and modulating key cytokines, including lowering pro-inflammatory IL-1β, IL-18 and TNF-α levels while normalizing anti-inflammatory IL-10 under inflammatory challenges. Importantly, nanoalgosomes exhibited excellent biocompatibility and no detectable cytotoxicity, underscoring their suitability for airway-targeted applications.

The antioxidant and immunomodulatory activities are likely attributed to the rich cargo of *T. chuii* bioactive molecules—such as carotenoids, polyphenols, and polyunsaturated fatty acids—combined with the intrinsic protective and delivery properties of EVs. Given their natural origin, scalability, and compatibility with aerosol-based administration, nanoalgosomes represent a promising therapeutic nanoplatform capable of restoring redox balance and immune homeostasis within the lung microenvironment.

Further investigations should delineate the molecular pathways underlying nanoalgosome-mediated bioactivity and confirm their therapeutic efficacy, pharmacokinetics, and safety in vivo using preclinical inhalation models. Collectively, these findings lay the groundwork for developing nanoalgosome-based inhalable therapeutics as a novel, biologically derived strategy for managing chronic inflammatory and oxidative lung diseases such as asthma, COPD, and pulmonary fibrosis.

## Acknowledgments

The authors would like to thank Nicolas Touzet for his assistance in establishing the microalgal cultures and the TFF setup, and Paolo Bergese for his insightful discussions on technical and conceptual aspects throughout the H2020 BOW project.

## Author contributions

**Wesam Darwish**: Methodology, Validation, Formal analysis, Investigation, Writing – Original Draft, Visualization. **Giorgia Adamo**: Methodology, Validation, Investigation, Writing – Review & Editing. **Mohammad Almasaleekh**: Validation, Investigation, Writing – Review & Editing. **Sabrina Picciotto**: Validation, Investigation, Writing – Review & Editing, Visualization. **Paola Gargano**: Validation, Investigation, Writing – Review & Editing. **Daniele P. Romancino**: Formal analysis, Writing – Review & Editing. **Samuele Raccosta**: Validation, Formal analysis, Investigation, Writing – Review & Editing. **Ralf Zimmermann**: Resources, Writing – Review & Editing, Supervision. **Mauro Manno**: Resources, Writing – Review & Editing, Supervision. **Antonella Bongiovanni**: Conceptualization, Methodology, Validation, Resources, Writing – Review & Editing, Supervision, Project administration, Funding acquisition. **Sebastiano Di Bucchianico**: Conceptualization, Methodology, Validation, Formal analysis, Writing – Review & Editing, Supervision, Project administration, Funding acquisition.

